# SIP: An Interchangeable Pipeline for scRNA-seq Data Processing

**DOI:** 10.1101/456772

**Authors:** Sijie Chen, Zheng Wei, Yang Chen, Kui Hua, Wei Zhang, Changyi Liu, Haoxiang Gao, Hao Sun, Zhenyi Wang, Qijin Yin, Shengquan Chen, Shaoming Song, Chen Feng, Hairong Lu, Rui Jiang, Xiaowo Wang, Jin Gu, Xuegong Zhang

## Abstract

Multiple steps of bioinformatics processing are needed to convert the raw scRNA-seq data to information that can be used in downstream analyses and in building cell atlases. Dozens of software packages have been developed and different labs tend to have different preferences on choices of the workflow. Such diversity can cause difficulties in future efforts of aggregating data from multiple labs, and also difficulties for new labs to start in this field. A few pipelines have been developed to help integrating multiple steps into a whole, but the fixed software architecture makes it hard for developers to add new features or exchange parts in the pipeline.

We presented SIP, a Single-cell Interchangeable Pipeline. It is a one-stop platform for the processing of scRNA-seq data from multiple platforms, and will also support for other types of data like scATAC-seq data. SIP utilizes container technology to solve the deployment dilemma when handling multiple packages and provides an easy-to-use interface for users to conduct the complicated multi-step process from raw data to final results with a single command. It also allows advanced users to assemble different versions of the pipeline by interchanging parts or adding new modules. SIP is available at https://github.com/XuegongLab/SIP under the GPL-3.0 license.

## 1 Introduction

Analysis pipelines for single-cell RNA sequencing (scRNA-seq) gain increasing attentions as scRNA-seq is becoming more and more important in biological studies (Wang and Navin, 2015; Hedlund and Deng, 2018; Wu *et al.*, 2017). ScRNA-seq is also one of the key technologies in the collaborative effort of building the human cell atlas (HCA)(Rozenblatt-Rosen *et al.*, 2017). Automated or semi-automated bioinformatics pipelines following standardized framework are one of the fundamental elements to support the effort.

To date, a group of pipeline and software packages have been developed for analysis of single-cell data. Some tools like simpleSingleCell (Lun *et al.*, 2016) guide users to manipulate data manually and require heavy programming skills. Some packages, such as Seurat (Butler *et al.*, 2018), Monocle 2 (Trapnell *et al.*, 2014; Qiu *et al.*, 2017), Scater (McCarthy *et al.*, 2017), ASAP (Gardeux *et al.*, 2017), and Granatum (Zhu *et al.*, 2017), mainly focused on the downstream analyzing steps after the gene expression quantification. However, the quantification of gene expression from raw data is not a trivial task. The alignment process often consumes substantial computational resources and the computation parameters correspond to different protocols can be very tricky. Therefore, a pipeline that starts from the raw data to downstream analysis is still highly demanded, especially for labs that just start to work in this field, and for labs that need to integrate their data with those from other labs. In addition, as the sequencing throughput is quickly getting higher, a scalable pipeline capable of utilizing the power of HPC (High-Performance Computing) clusters is in urgent need for labs and consortiums that generate massive single-cell data.

To meet these needs, we developed a Single-cell Interchangeable Pipeline called SIP. It is an open-source pipeline for processing from raw data to final analyses with standard framework and interchangeable components. SIP contains seven main modules, including Read QC and preprocessing, (Quasi-)alignment and transcript assembly, Gene-level abundance counting, Cluster finding, Trajectory analysis, and DEG analysis. Each module contains multiple encapsulated tools. The tools can be easily replaced with alternatives, and new tools and/or modules can also be added.

SIP is constructed with Nextflow (DI Tommaso *et al.*, 2017), which seamlessly take over outputs and inputs of the upstream and downstream modules, significantly alleviating the data manipulation efforts for users. SIP adopts the container technology to deploy the pipeline. Several useful features including e-mail notification and HPC cluster support make it more convenient to process massive amount of data using SIP. Besides, it is easy to extend SIP’s function as Nextflow has a large developer community.

Excellent computational tools constantly appear in the field of single-cell data analysis. To keep pace with the development of new tools, a good pipeline should have an open software architecture for changes and be endorsed by a development community so that new tools can be integrated efficiently. The framework used in SIP allows very easy maintaining and updating the pipeline, and more potential Nextflow developers can join the improvement of SIP in a collaborative manner.

## 2 Methods

SIP prepares a curated list of single-cell RNA-seq data processing tools. The input and output of each tool are parsed by the Nextflow framework and transferred automatically. As shown in Fig. 1, users can choose the interchangeable parts for a specific step. With different combination of components, workflows of different steps are established to process the data. Users can easily explore proper tool sets for different datasets, and make a performance comparison on different datasets.

**Fig. 1.**
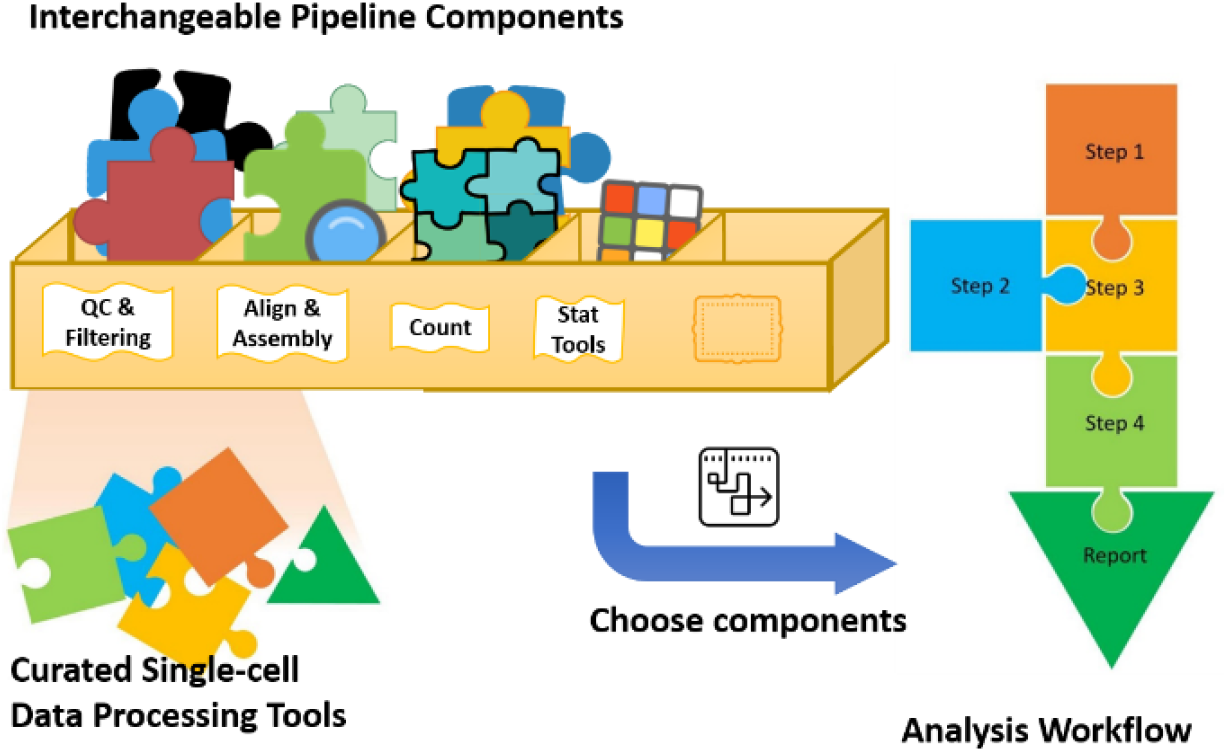
Concept map of SIP. A group of software and packages are evaluated and selected to build up interchangeable parts in the pipeline toolbox. Users can freely choose the tools to process the data without concerning the tedious intermediate data manipulation operations.

### 2.1 The SIP workflow

#### 2.1.0 Feed raw sequencing data to pipeline

SIP accepts raw sequencing data (fastq files) as input. Read files should be named after identifiers of the cells and placed in the cell home folder. The pipeline detects all files in the folder and group the files by name. The layout (single-end, paired-end) of sequencing library will be automatically determined and all layout-associated parameters will be set by the SIP. Please check the online documentation for more details of read arrangements.

#### 2.1.1 Read quality control and preprocessing

Various quality defects (e.g. adapter sequences, low-quality bases, poly-T ends etc.) may exist in the raw sequencing reads. Directly processing the raw reads would lower down the quality of results too (Del Fabbro et al., 2013), hence it is necessary to check the quality of reads before further steps and filter out defective reads and produce clean reads for succeeding steps. SIP integrates serveral tools for read quality checking and filtering, including FastQC (Andrews, 2010), fastp (Chen et al., 2018), AfterQC (Chen et al., 2017), and applies the quality control and preprocessing step to each sample or cell. Separated results for each cell are further summarized with MultiQC (Ewels et al., 2016).

#### 2.1.2 (Quasi-)Alignment of reads and transcript assembly

Alignment (or quasi-alignment) or short sequencing reads is the essential step in the reference-based transcriptome studies. SIP automatically detects the read layout and the “strandness” of the RNA-seq experiment with RSeQC (Wang et al., 2012) and set the corresponding parameters for users in the (quasi-)alignment steps.

SIP by default uses a quasi-alignment tool, salmon (Patro et al., 2017), for transcript quantification because it is much faster than alignment-based tools. For alignment-based transcript assembly, we include several classical packages in SIP: STAR (Dobin et al., 2013) and Hisat2 (Kim et al., 2015, 2017) for read alignment, StringTie (Pertea et al., 2015) for transcript assembly. Like the previous Read QC and Preprocessing step, the (quasi-)alignment qualities (e.g. read mapping rate, aligned read count, fragment length distribution) are also summarized and reported by MultiQC.

#### 2.1.3 Gene-level abundance counting

For alignment-based quantification, the generated Sequence Alignment Map files (e.g. SAM, BAM files) can be either counted with featureCounts (Liao et al., 2014) or HTSeq (Anders et al., 2015) at read level, or counted based on normalized results (TPM, RPKM, or FPKM) produced by StringTie.

For the quasi-alignment method, abundance levels of transcripts are directly estimated without explicitly assembling the transcriptome. Normalized results or read counts are supplied by salmon in SIP.

As accurate quantification of transcriptional isoforms at single-cell level is still challenging (Vu et al., 2018; Arzalluz-Luqueángeles and Conesa, 2018), SIP currently imports all cells’ transcript-level quantification results and merge them into gene-level expression matrix where each column depicts an expression profile of a single cell with the help of the R package tximport (Soneson et al., 2016).

#### 2.1.4 Post-quantification downstream analysis

SIP can take the gene-level expression matrix generated by the previous abundance counting step, and continue to conduct downstream analyses like clustering, identifying differentially expressed genes, and building cell trajectories, etc. A number of packages have been developed to accomplish such statistical analysis. SIP has currently integrated Seurat, Monocle 2, and Scater.

### 2.2 Other useful features

By utilizing the advantages of the Nextflow framework, SIP has the following useful features:

- **User-friendly reports**: Important intermediate results and summarized final results are well-organized in the output directory along with multiple user-friendly HTML and PDF graphical reports.
- **Cluster environment**: Nextflow supports the deployment on cluster environment, therefore the pipeline is capable of processing very big data with the slurm HPC cluster management system.
- **Parallel processing**: Processes can be computed in a parallel manner, maximizing the usage of computational resources.
- **Breakpoint resume**: SIP caches accomplished work with Nextflow. Finished processes will not be wasted when a very long pipeline is interrupted in the middle.
- **Container support**: SIP provides Docker images with all software and packages installed inside. Users can plug and play the pipeline directly without pre-install the numerous dependencies.
- **E-mail notification**: SIP can be configured to send information automatically when a certain process or the whole workflow is finished.
- **Developer community**: Nextflow has a large developer community. This will help the future development and extension of SIP.

## 3 Results

We applied SIP on several single-cell RNA-seq datasets to test the usability and performance of the pipeline. We selected a subset of the single-cell dataset generated by SMART-seq with accession number GSE38495 (Ramsköld et al., 2012). The subset contains 8 hESCs (human embryonic stem cells), and 19 LNCaP cells (androgen-sensitive human prostate adenocarcinoma cells). The whole analysis workflow is finished in 46 minutes on a workstation with 2 Intel^®^ Xeon^®^ E5-2670 CPU and 96 GB memory. The real memory requirement for the package depends on the particular set of modules uses choose in the pipeline. The actually memory consumption in this example experiment is <40GB, and we have also checked that SIP can be installed and ran on personal computers or laptops with memory as low as only 8GB.

The raw sequencing reads are checked and filtered with fastp using the “--readqc=fastp” option. Pre-processed clean reads are quantified in quasi-alignment mode with salmon using the “--quant=salmon” option. Fig. 2 (a) shows snapshots of read quality control results and quasi-alignment results. In general, the read qualities of 27 samples are good enough to perform succeeding steps: 97.5%~99.5% of the reads passes the read filtering; adapter sequences are detected in 12 out of the 27 samples, and the percentage of adapter-trimmed reads are ≤ 1.0%. More details about GC-content, sequence quality, duplication rates, N contents, etc. are also suppled in the report.

After quantification, the transcript-level expression profiles of multiple cells are automatically transformed into gene-level and then merged into a matrix of all cells to feed the downstream statistical analysis. Clustering identification and differentially expressed gene identification are conducted using Seurat with the “--using_seurat” option for this dataset. We used a typical Seurat workflow:

1) Initialize a Seurat object;
2) Filter cells based on built-in metrics;
3) Normalize the data;
4) Select variable genes across cells;
5) Removing confounding factors by regression.
6) Linear dimensionality reduction.
7) t-SNE visualization
8) PCA-based clustering
9) Identify differentially expressed genes

**Fig. 2.**
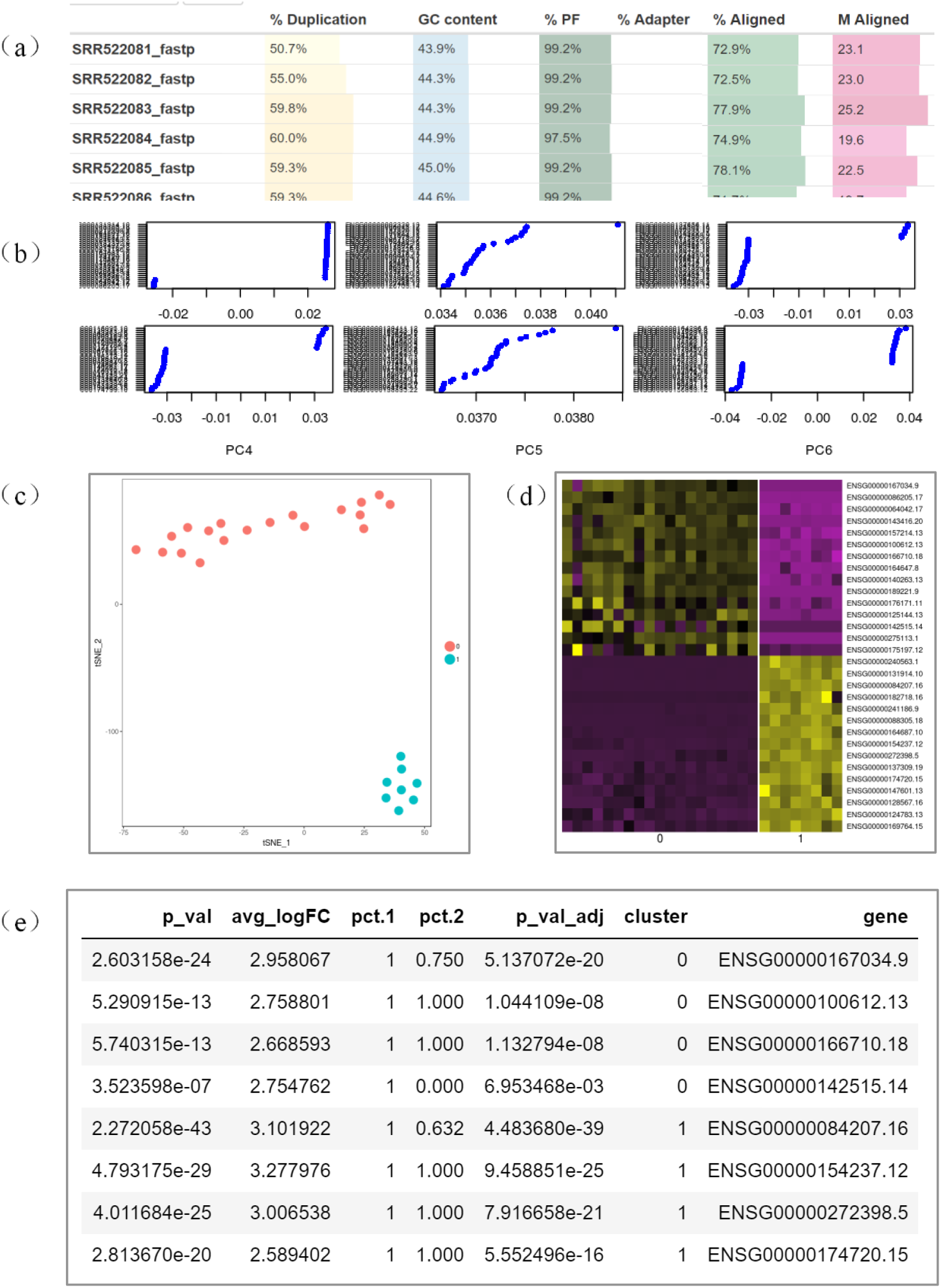
Selected results of experiment 1. (a) read quality check report and quasi-alignment report; (b) Visualizations of PCA’s first 6 components; (c) t-SNE plot of the 2 types of cells; (d) Heatmap of the cells across part of selected marker genes; (e) Selected differentially expressed marker genes.

Fig. 2 (b) visualizes the result of PCA linear dimensionality reduction with SIP. The first 6 principle components are listed, where x-axis represents the PC’s value, and y-axis represents the corresponding feature genes. Fig. 2 (c) is the t-SNE plot of the PCA-transformed data reported by SIP. The PCA-based clustering algorithm obtains two cluster and they exactly correspond to the two real cell types (19 LNCap cells and 8 hESCs). The colors in the t-SNE plot are inferred from cluster information automatically by SIP. The t-SNE is derived with perplexity parameter = 5 and first two PCA dimension. The clustering algorithm used the first 10 principle components and a resolution parameter = 1.04.

Differentially expressed marker genes are then identified between the two computed groups. A heatmap with an obvious pattern shows top 15 genes with high fold changes in Fig. 2 (d). Fig. 2 (e) lists the most significant differentially expressed marker genes in two cell types.

## 4 Conclusion

We have developed a one-stop pipeline SIP for processing and analyzing single-cell RNA-seq data from the raw fastq data to downstream analyses and automatic report generation. The pipeline is interchangeable and expansible, and can be scaled up to be ran on HPC clusters. The current version of SIP supports data generated from all major RNA-seq platforms. And the future upgrades will soon support other types of single-cell sequencing data such as scATAC-seq data. With this handy pipeline, biological labs that generating single-cell sequencing data can be freed from the tedious and challenging labors of installing and running multiple software packages and setting computer environments to make everything compatible. This allows them to better focus on the scientific questions to be answered from the data. Besides, the pipeline also allows biological labs to be able to share bioinformatics pipelines including detailed parameter settings with other labs, which makes it easier for integrating data from multiple labs in big collaborative projects such as the Human Cell Atlas (HCA) program. SIP is available at https://github.com/chansigit/SIP under the GPL-3.0 license.

## Acknowledgements

We thank Zhun Miao, Xiangyu Li, Dongfang Wang and Dongyang Luo for their helpful discussions.

Conflict of Interest: none declared.

